# Targeting CXCR4 with [^212^Pb/^203^Pb]-Pentixather Significantly Increases Overall Survival in Small Cell Lung Cancer

**DOI:** 10.1101/2024.09.06.611663

**Authors:** Keegan A. Christensen, Melissa A. Fath, Jordan T. Ewald, Claudia Robles-Planells, Stephen A. Graves, Spenser S. Johnson, Zeb R. Zacharias, Jon C.D. Houtman, M Sue O’Dorisio, Michael K. Schultz, Bryan G. Allen, Muhammad Furqan, Yusuf Menda, Dijie Liu, Douglas R. Spitz

## Abstract

**Introduction:** Small cell lung cancer (SCLC) has a 7% 5-year overall survival. C-X-C chemokine receptor 4 (CXCR4), an attractive target for theranostic agents, is highly expressed in SCLCs, and can be targeted with pentixather using the theranostic pair ^212^Pb/^203^Pb. The hypothesis that [^212^Pb/^203^Pb]-pentixather can be used safely and effectively for imaging and therapy in SCLC in xenograft models was tested.

**Results:** SPECT/CT imaging and biodistribution studies of tumor bearing mice injected with [^203^Pb]-pentixather demonstrated CXCR4-dependent uptake in tumors and accumulation of radioligand in kidneys and livers. Dosimetry calculations estimated [^212^Pb]-pentixather uptake in tumor and normal tissue. [^212^Pb]-pentixather treatment (37-111 kBq/g) of SCLC xenografts (DMS273 and H69AR) significantly prolonged survival and delayed tumor growth. When NSG mice grafted with human hCD34+ bone marrow were treated with [^212^Pb]-pentixather (37-111 kBq/g), significant cytopenias were observed in peripheral blood complete blood counts (CBCs) at 13-18 days post treatment which resolved by day 28-31. Flow cytometry of bone marrow hematopeotic stem cells in these animals at day 28-31 demonstrated a significantly reduced frequency of the human hematopoietic marker CD45 (hCD45+) and reconstitution of the bone marrow with murine CD45+ (mCD45+) lineages.

**Conclusions:** [^203^Pb]-pentixather can be used to image CXCR4 expressing SCLC xenografts and treatment with alpha emitter [^212^Pb]-pentixather significantly prolongs SCLC xenograft median overall survival. Significantly greater mCD45+ bone marrow repopulation was detected in NSG mice engrafted with human bone marrow 28-31 days following [^212^Pb]-pentixather treatment, relative to hCD45+ bone marrow.

## Introduction

Small cell lung cancer (SCLC) conveys a bleak prognosis with a 5-year survival of only 7% ^1^. Therefore, it is crucial to develop new therapies for SCLC. Peptide receptor radionuclide therapy (PRRT) involves the targeted delivery of radionuclides to receptors expressed on the surface of tumor cells. C-X-C chemokine receptor 4 (CXCR4) is normally involved in the mobilization of hematopoietic stem cells but is also highly expressed in some SCLC. CXCR4 expression has been shown to be significantly associated with mortality, with high expressers 2.5 times more likely to die than low expressors ^2,3^. Pentixather, a ligand which exhibits high affinity for human CXCR4, can be radiolabeled for use in both therapy and imaging of CXCR4 positive tumors ^4,5^.

Alpha particle emitting radionuclides have reinvigorated interest in PRRT as they offer high linear energy transfer (LET) with decreased off-site tissue damage compared to beta emitters. The alpha emitter ^212^Pb and its elementally-matched imaging surrogate ^203^Pb have been studied in the context of multiple cancers and targeting vectors ^6^. Alpha particles produce dense ionization tracks that result in highly lethal DNA damage when nuclear traversal occurs, relative to low LET radiation ^7^.

The current study tested the hypothesis that SCLC xenograft tumors can be effectively imaged and treated with the theranostic pair [^203/212^Pb]-pentixather ([^203/212^Pb]-pent). We utilized [^203^Pb]-pent single photon emission computed tomography (SPECT/CT) in mice bearing human SCLC tumors with increasing levels of CXCR4 to demonstrate that tumor specificity correlates to the receptor expression. Dosimetry calculations were performed, which correlated with findings of biodistribution studies. [^212^Pb]-pent treatment of SCLC xenografts resulted in prolonged survival with minimal off target kidney, liver, and bone marrow damage. Humanized hCD34+ NSG mice carrying both human and murine hematopoietic lineages were utilized to investigate the hematologic toxicity of [^212^Pb]-pent at therapeutic doses 28-31 days after treatment and demonstrated significant reductions in the frequency of hCD45+ bone marrow compared to mouse bone marrow cells. However, a surviving population of viable human cells was detectable in all animals, and a minority of animals retained considerable human hematopoiesis. The results support continuing investigation of [^203/212^Pb]-pent as a theranostic tool in SCLC.

## Methods

### Mice

Animal experiments were approved by the IACUC committee of the University of Iowa and conformed to the guidelines established by NIH. Four-to six-week-old female athymic nu/nu mice were obtained from Envigo for therapy and imaging studies. Additional four-to six-week-old female athymic nu/nu mice were purchased from Charles River for the [^212^Pb]-pent therapy study. Humanized hCD34+ mice were purchased from Jackson Laboratory (Jackson 705557 Hu-NSG-CD34) and generated from NOD/SCID/IL2rγ^null^ mice injected with human hCD34+ hematopoietic stem cells from 6 donors **Supplemental Table 2** ^8, 9^.

### Cell culture conditions

SCLC lines DMS53 and H69AR and non-small cell lung cancer cell line (NSCLC) H292 were obtained from the American Type Culture Collection and DMS273 cells from ECSCC through Sigma-Aldrich. Cell lines were regularly tested for mycoplasma, and grown in RPMI1640 medium with 10% FBS (Hyclone). Cells were maintained at 37°C with 5% CO_2_ and were utilized within 20 passages after purchase.

### ^203^Pb/^212^Pb radiolabeling for SPECT imaging and *in vivo* studies

^203^Pb chloride (obtained from Medical Isotope and Cyclotron Facility, University of Alberta, Canada) or ^212^Pb chloride (eluted with 3 ml of 2 M HCl from a radium-224 (^224^Ra)/^212^Pb generator, Perspective Therapeutics) was purified as described ^10^. The Pb isotopes (pH=6 in NaOAc buffer; 1 mL) were added to 1 M pH=4 NaOAc buffer containing pentixather (PentixaPharm) with ascorbate (1 mg/ml, pH 5.0). Post radiolabeling at 85°C for 25 min, the mixture was purified by a C-18 reverse-phase cartridge (Strata-X Solid Phase Extraction, SPE; Phenomenex) and eluted in 50% ethanol saline solution. The radiochemical yield was 86.8% with molar activity of 63.7 MBq/nmol and 70% with molar activity of 7.4-10.0 MBq/nmol for [^203^Pb]-pent and [^212^Pb]-pent, respectively.

### Micro-SPECT/CT imaging and biodistribution

Six nu/nu mice were implanted with 1-10 million SCLC cell lines with DMS53 (left shoulder), H69AR (right shoulder), and DMS273 (right flank) with H292 NSCLC negative controls (left flank). Two to three weeks post inoculation, when the tumors reached 3-10 mm in diameter, three mice were administered [^203^Pb]-pent via tail vein (approximately 11.1 MBq each) and imaged at 0.5, 4, and 24 h post injection using a micro-SPECT/CT imaging instrument (Siemens Inveon). Mouse organs and tumors were harvested 30 h post-injection, weighed, and counted using Packard Cobra II Auto-Gamma Counter. Radioactivity was presented as % injected dose per gram of tissues (%ID/g) with decay correction.

### Dosimetry Calculations

The absorbed dose per administered activity was estimated for a theoretical injection of [^212^Pb]-pent based on the time-ordered [^203^Pb]-pent SPECT/CT images obtained from scans in **Figure 1A**. Scans were calibrated to provide image units of Bq/voxel based upon a known quantity of ^203^Pb in a spherical sensitivity calibration phantom. Quantitative SPECT/CT images were segmented in MIM Software (v 7.3.2) to determine organ (kidney and liver) and tumor time-activity curves. Partial-volume correction was performed by region of interest expansion with manual adjustment to limit overlap with adjacent structures. Organ and tumor time-activity curves were multiplied the ratio of ^212^Pb to ^203^Pb decay constants 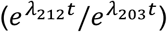 to simulate the appropriate physical decay of [^212^Pb]-pent. Resulting time activity curves were integrated using the trapezoidal method with physical decay following the 24-hour imaging timepoint (when <5% of the injected [^212^Pb]-pent would remain in the body) to generate data in **Figure 1B**.

**Figure 1.**
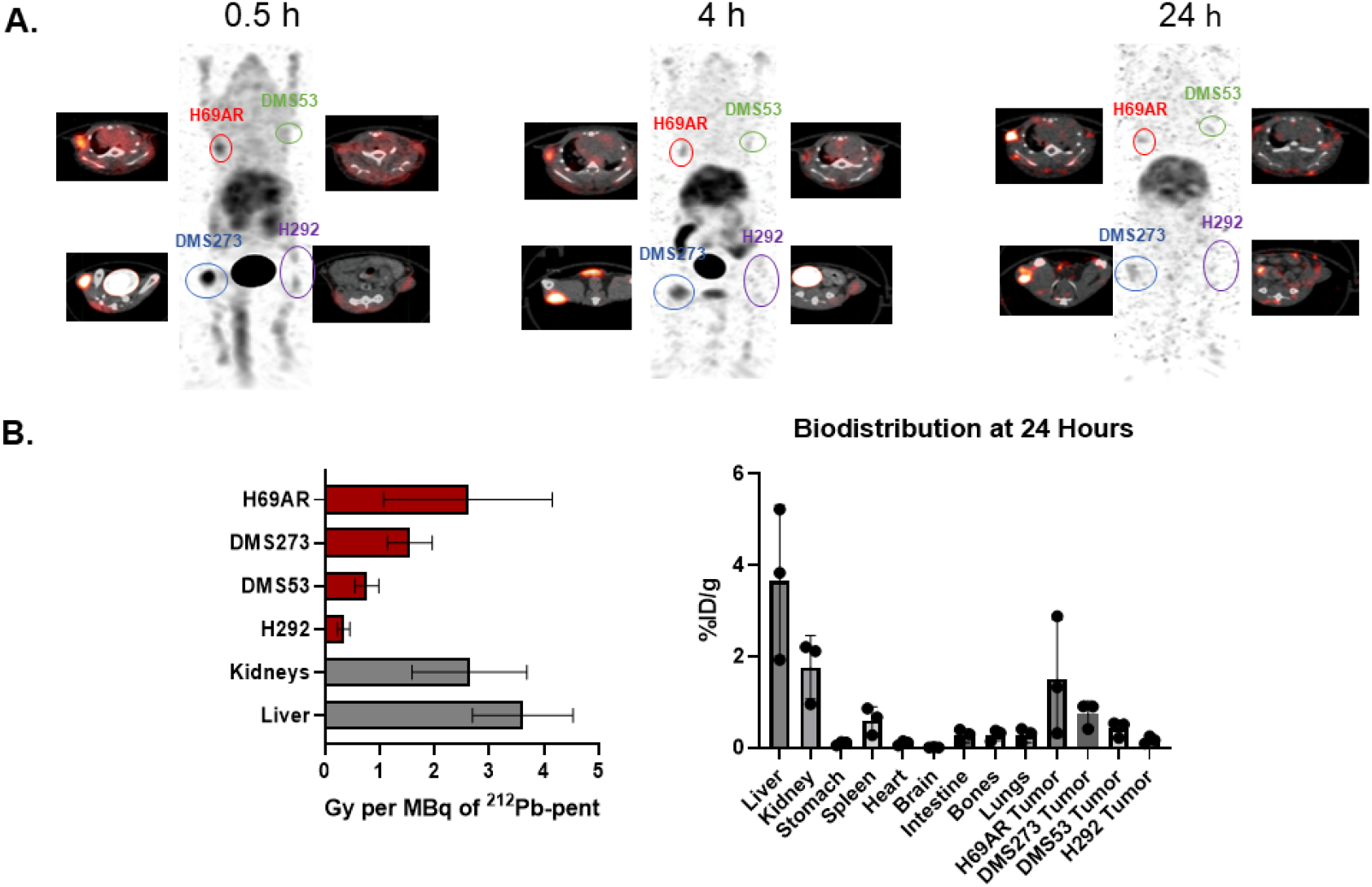
Preclinical [^203^Pb]-pent Spect/CT imaging in nude mouse xenografts estimates tumor and organ uptake and allows for dosimetry estimates. **A)** Four human lung cancer xenografts were grown on 3 nude mice and then injected with 11.1 MBq [^203^Pb]-pent and SPECT/CT image performed 0.5, 4 and 24 hours (h) post injection. **B)** Dosimetry of detected activity was used to determine the average absorbed doses in liver, kidneys, H292 NSCLC negative control xenografts, and DMS53, DMS273, H69AR SCLC xenografts. Biodistribution of ^203^Pb was determined at 30 hours as percent injected dose per gram of tissue (%ID/g) as measured via γ-counter.

### [^212^Pb]-pent efficacy study in tumor-bearing mice

For efficacy studies, tumor cells (1 × 10^6^ DMS273 cells) were injected into nu/nu mice flanks and when tumor volume reached ∼30 mm^3^, by [Vernier calipers Volume = (Length X Width^2^)/2]), mice were randomly assigned to groups. Mice obtained from two vendors, were evenly split between treatment groups. Intraperitoneal injection of saline/arginine/lysine was given, before and after IV injection of [^212^Pb]-pent, as described ^3^. Tumor volumes, body weights, and adverse events were recorded daily by a blinded investigator. When tumor volume exceeded 1000 mm^3^ for two measurements animals were euthanized unless otherwise noted.

### Bone Marrow Analysis shown in Figure 2D

Whole bone marrow cells were obtained by flushing one femur with PBS 2% FBS followed by erythrocyte elimination with ACK Lysing Buffer (10-548E, BioWhittaker, Lonza). The remaining cells were filtered through 40 μm pore size cell strainer and counted. 3×10^6^ cells were stained with Zombie Aqua (423102, BioLegend) for 20 min at room temperature for live/dead cells discrimination, followed by a 30 min surface staining on ice using PBS 1% BSA as staining buffer with the following antibodies: Lineage negative cocktail, Lin—APC (BD Bioscience), Sca1-PerCP/Cy5 (Clone D7, BioLegend), cKit-APC/Cy7 (Clone 2B8, BioLegend), CD135-PE (Clone A2F10, BioLegend), CD48-FITC (Clone HM48-1, BioLegend) and CD150-PE/Cy7 (Clone TC15-12F12.2, BioLegend). Cells were immediately analyzed using a Cytek® Aurora flow cytometer (Cytek Biosciences Inc.) with five lasers (355, 405, 488, 561, and 640nm) and 64 detectors. The long-term hematopoietic stem cell (LT-HSC) population was defined as Live LincKit^+^ Sca1^+^ CD135^-^ CD48^-^ CD150^+^ cells using FlowJo Software v10.8.1 (Treestar, Ashland, OR). Gating scheme shown in Supplemental Figure 3.

**Figure 2.**
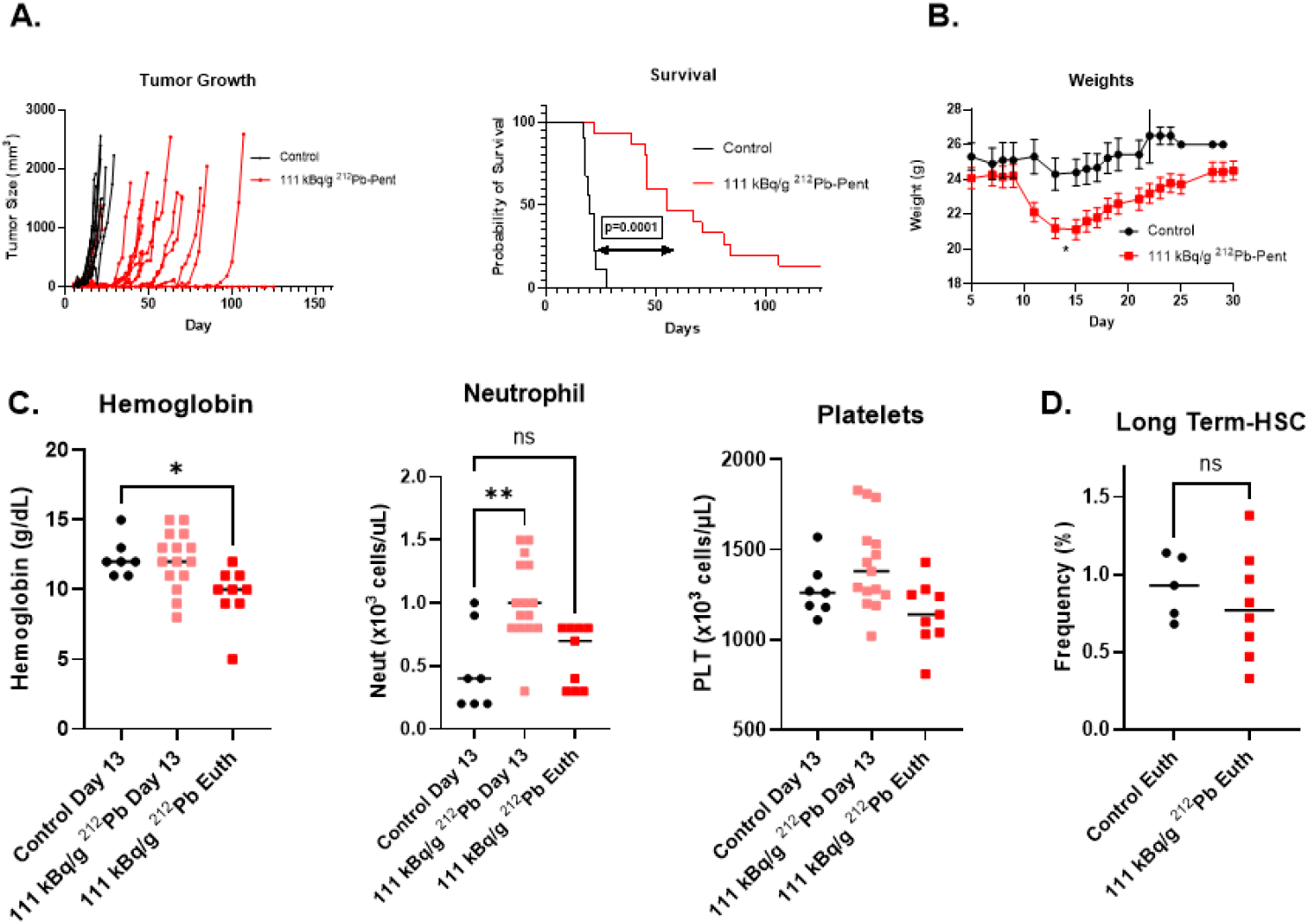
[^212^Pb]-pent therapy at the 111 kBq/g significantly prolonged survival and delayed tumor growth with tolerable bone marrow toxicity in nude mice bearing SCLC xenografts. 111 kBq/g of body weight (n=15) or equivalent volume of saline control (n=10) was injected into nude mice bearing DMS273 xenografts. **A)** Xenograft tumor volume was measured daily and each mouse graphed. Mice were euthanized when tumors exceeded 1000 mm^3^ for two measurements. Kaplan-Meier survival analysis demonstrated a significant difference comparing treated vs controls. **B)** Animal weights were recorded daily and significant weight loss was observed in treatment days 11-22 as determined by multiple unpaired t-tests (p<0.05). **C)** Mice were bled via mandibular vein at day 13 and euthanasia and CBCs were performed. Hemoglobin, neutrophil and platelets were graphed (p<0.05, one way ANOVA, Dunnett’s multiple comparisons) **D)** The long term hemopoietic stem cells from a bone marrow flush were stained and subjected to flow cytometry (Gating shown in **Supplemental Figure 3**) was not significantly different with treatment as determined by unpaired t-test.

**Figure 3.**
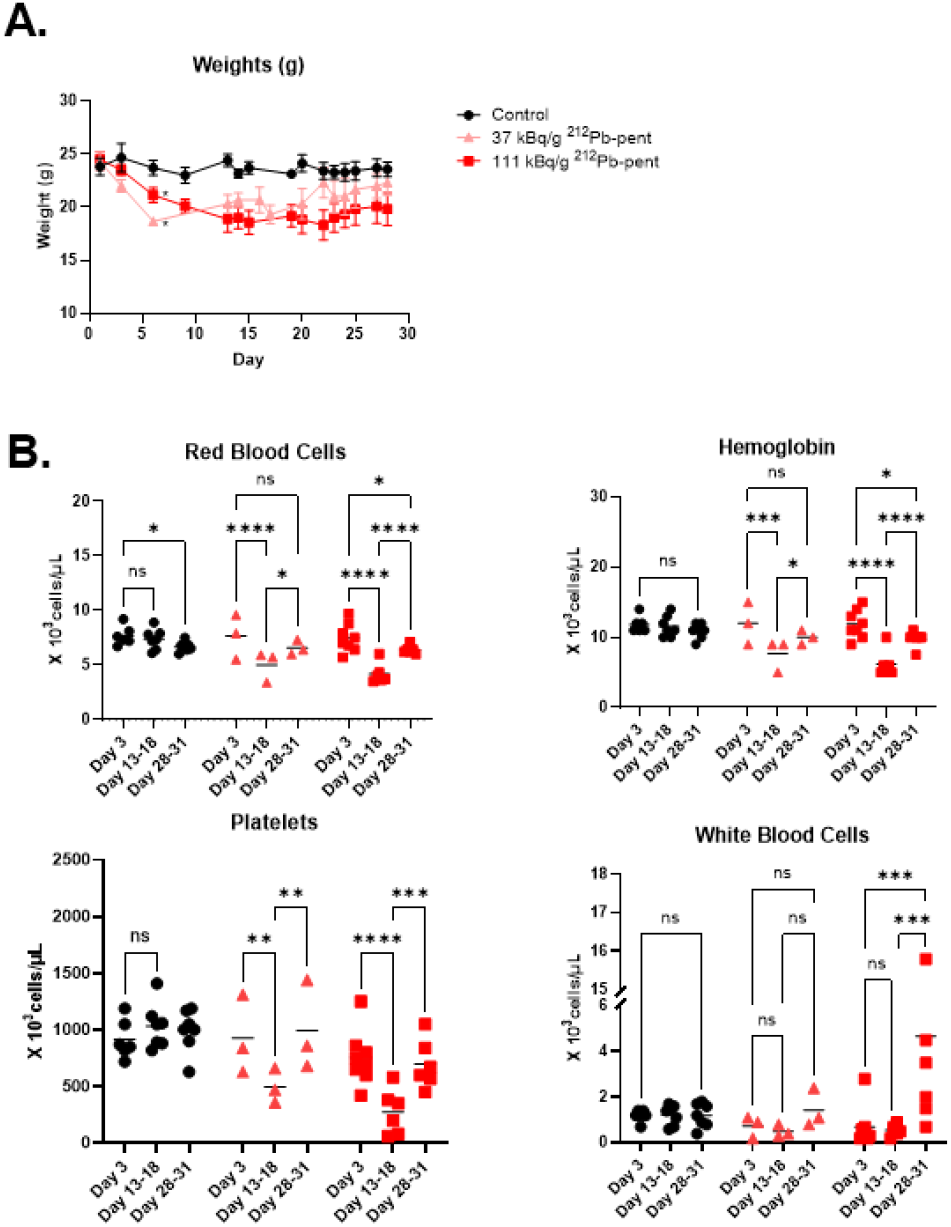
[^212^Pb]-pent causes transient cytopenias in humanized hCD34+ mice. NOD/SCID/IL2rγ^null^ mice with functional hCD34+ bone marrow were injected with 37 kBq/g [^212^Pb]-pent (n=3), 111 kBq/g [^212^Pb]-pent (n=9) or saline control (n=8). **A)** Animal were weighed daily and averages graphed with significance determined by multiple unpaired t-tests (p<0.05 day 6-24). **B)** Animals were bled via submandibular vein on day 3, day 13-18, and day 28-31 and CBCs were performed and graphed (p<0.05, one way ANOVA, Dunnett’s multiple comparisons).

### ^212^Pb dose calibration

To standardize ^212^Pb radiopharmaceutical doses, we obtained a ^232^U source (Eckert & Ziegler; 3.391 kBq) which maintained radioactive equilibrium of ^212^Pb with its decay daughters. On the day of an experiment, the counting efficiency (62.0–63.7%) of ^212^Pb-specific peak (239 keV) was determined for 2 mins using a sodium iodide scintillation detector, taking into account the intensity (43.6%) of the 239 keV photons. Counts per sec (CPS) ^212^Pb at 239 keV CPS were converted to ^212^Pb activity using the pre-determined efficiency and intensity of the 239 keV photons. These calculated values were in agreement with theoretical values (<10%).

### Complete blood cell counts

A lancet was used to draw blood from the submandibular vein of mice. Blood was diluted 1:10 in PBS and CBC were processed in a Siemens ADVIA 120 Hematology Analyzer.

### Flow cytometry of human hCD34+ NSG mice

Bone marrow cells were obtained by flushing two femurs of each NSG mouse with 2% FBS in PBS followed by erythrocyte elimination with ACK Lysing Buffer (10-548E, BioWhittaker, Lonza). Cells filtered through 40 μm strainer were counted and 1×10^6^ cells/well were stained with Live/Dead Blue (Thermo) for 15 min. Cells were subsequently stained with antibodies in FACS buffer (2% FBS, 0.2% sodium azide in PBS) supplemented with Brilliant Stain buffer, blocking serum (rat, hamster, and mouse), and CellBlox^TM^ Blocking Buffer for 30 min. The surface marker antibodies used were: LIVE/DEAD Blue (Thermofisher), anti-human CD16 BUV496 (clone: 3G8, BD Biosciences), anti-human CD14 BUV563 (clone: 63D3.rMAb, BD Biosciences), anti-human CD56 BUV737 (clone: NCAM16.2, BD Biosciences), anti-human CD3 BV510 (clone: SK7, Biolegend), anti-human CD45 PerCP (clone: 2D1, Biolegend), anti-human CD19 NovaFluor Red 700 (clone: HIB19, eBioscience/Thermofischer), and anti-human CD11c PE-Cy7 (Clone: 3.9, Invitrogen). Cells were washed with FACS buffers and fixed with BD FACS Lysing solution for 10 min. Following fixation, cells were washed, and stored in PBS at 4°C until analysis. Single stained cells were used as compensation controls. Gating strategies shown in Supplemental Figure 5. Data were collected on a Cytek® Aurora and analyzed with SpectroFlo and FlowJo software.

## Results

### Tumor dose correlates to CXCR4 expression using SPECT/CT imaging with [^203^Pb]-pent

SCLC cell lines were selected based on CXCR4 expression (see **Supplemental Table 1**) as measured by flow cytometry previously published ^3^. H292 (no expression), DMS53 (low expression), DMS273 (medium expression), and H69AR (high expression) cells were subcutaneously injected into both flanks and shoulders. Tumor xenografts varied from 3.2-13.2 mm in diameter and mice (N=3) were injected with 11.1 MBq of [^203^Pb]-pent via tail vein followed by SPECT/CT at 30 minutes, 4 hours, and 24 hours. Imaging shows high uptake in H69AR and DMS273 tumors and little to no uptake in the DMS53 and H292 tumors (**Figure 1A)**. Dosimetry estimates of [^212^Pb]-pent obtained from time-ordered SPECT/CT images in tumors, livers and kidneys (**Figure 1AB)** demonstrated averaged activity of 0.34 Gy/MBq in H292 xenografts, 0.77 Gy/MBq in DMS53, 1.5 Gy/MBq in DMS273, and 2.6 Gy/MBq in H69AR, consistent with CXCR4 expression. Furthermore, accumulation of [^203^Pb]-pent and ^212^Pb dose estimates in kidneys (2.6 Gy/MBq) indicated renal excretion and also retention in liver (3.6 Gy/MBq) at 24 hours (**Figure 1AB**). Residual radiation at 30 hours using a γ-counter expressed as initial dose per gram of tissue (%ID/g) (**Figure 1B, right panel)** accurately reflected the findings from SPECT/CT imaging with activity in the bone marrow being less than 0.5% ID/g measured in all three animals.These data support the utility of [^203^Pb]-pent as both a probe for CXCR4 expression and ^212^Pb-dosimetry calculations.

### 111 kBq/g [^212^Pb]-pent results in prolonged survival in SCLC xenografts

A single dose of 111 kBq/g [^212^Pb]-pent injected into nude mice bearing DMS273 SCLC xenografts (15 treatment and 10 controls) led to a highly significant delay in tumor growth rate (p<0.01) and increased median overall survival (**Figure 2A**, p<0.01) demonstrating anti-tumor efficacy with tolerable weight loss (**Figure 2B**). The experiment was repeated with DMS273 xenografts with [^212^Pb]-pent (111 kBq/g and 37 kBq/g), (9-10 mice per group) with tolerable weight loss as well as significant delays in tumor growth (p<0.05) and increased median overall survival (p<0.05) (**Supplemental Figure 1 A-F**). To verify efficacy in a multi-drug resistant SCLC cell line expressing CXCR4 (**Supplemental Table 1**) N=4 H69AR xenografts were treated with 55.5 kBq/g [^212^Pb]-pent, again causing a significant growth delay and increased median survival, with 3 of 4 treated mice showing no evidence of disease (**Supplemental Figure 2**). Taken together, these results suggest that [^212^Pb]-pent has the potential to be a highly effective treatment for CXCR4 expressing SCLC.

### 111 kBq/g [^212^Pb]-pent causes minimal normal tissue injury in nude mice

Randomly selected mice from groups shown in **Figure 2A** were bled 13 days post treatment as well as prior to euthanasia. CBCs demonstrated hemoglobin levels were reduced ∼15% at the time of euthanasia, a transient increase in neutrophils was observed on day 13 that resolved by euthanasia, and no significant changes in platelet counts were detected (**Figure 2C)**. After euthanasia, whole bone marrow was flushed from two tibias and one femur of each animal sampled and flow cytometry was performed to assess mouse hematopoietic stem cells using the gating strategy described in **Supplemental Methods Figure 3**. No changes in the frequency of long-term hematopoietic stem cells (LT-HSC) was noted with [^212^Pb]-pent treatment (**Figure 2D**). These data demonstrate treatment with [^212^Pb]-pent was well-tolerated by the hematopeotic system of xenografted mice at an effective therapeutic dose, although effects on human bone marrow cannot be extrapolated due to pentixather’s higher affinity for the human CXCR4 isoform ^4^.

Given the kidney and liver uptake seen on SPECT/CT ^203^Pb, 5 randomly selected mice from the experiment shown in **Supplementary Figure 1FA-C** were analyzed at euthanasia and serum chemistries for liver and kidney injury markers were performed. Two mice in the treatment group exhibited elevated levels of ALT, a serum marker of liver damage (**Supplemental Figure 4**). There was no observed difference between treatment and control groups in serum protein, alkaline phosphatase, or bilirubin (**Supplemental Figures 4A, C, D**). Blood urea nitrogen (BUN) and serum creatinine did not differ between control and treatment groups (**Supplemental Figure 4E, F**). These findings support the hypothesis that [^212^Pb]-pent caused minor discernable damage to kidneys or livers at therapeutic doses.

**Figure 4.**
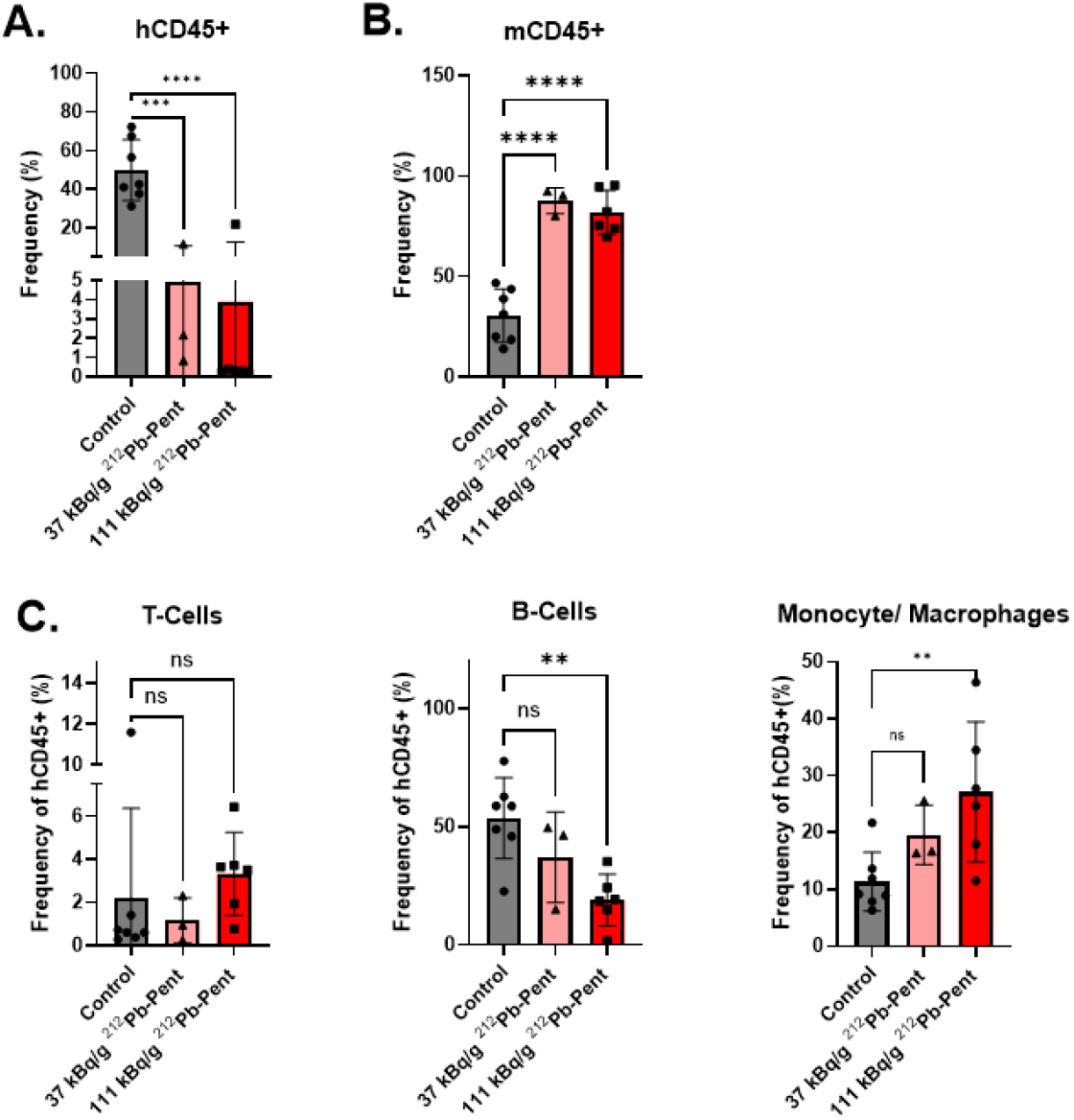
Treatment of humanized hCD34+ mice with 111 kBq/g [^212^Pb]-pent resulted in reduced frequency of human cell types and a repopulation with mouse bone marrow. Humanized hCD34+ mice treated with 37 kBq/g [^212^Pb]-pent (n=3), 111 kBq/g [^212^Pb]-pent (n=9) or saline control (n=8) were sacrificed on day 28-31 and bone marrow was harvested. Marrow was stained for human and mouse hematopoietic lineages and flow cytometry was performed. (Gating shown in **Supplemental Figure 5**) **A)** The percentage of viable hCD45+ cells in the treatment cohorts compared to control. **B)** The percentage of viable mCD45+ cells in [^212^Pb]-pent treated animals compared to controls. **C)** Human T-cell, B-cell, and monocyte/macrophage frequency among hCD45+ cells in the [^212^Pb]-pent treated mice compared to control. Significance as determined by one way ANOVA with Fisher’s least significant difference (p<0.05).

### [^212^Pb]-pent treatment in humanized hCD34+ mice results in significant myelotoxicity with viable human cells detectable in all animals

Bone marrow toxicity is a concern for CXCR4 targeted therapy due to high expression of CXCR4 on HSCs. Since pentixather is known to exhibit greater affinity for the human isoform of CXCR4 compared to the murine isoform ^4^, experiments were conducted utilizing humanized hCD34+ mice. Nine hCD34+ mice were randomly assigned to receive saline control (N=3), 37 kBq/g [^212^Pb]-pent (N=3), or 111 kBq/g [^212^Pb]-pent (N=3) via tail vein injection. The experiment was then repeated with an additional 9 mice (N=4 control, N=5 111 kBq/g [^212^Pb]-pent) and results were pooled. The vendor quantified the expression of both human and mouse lineages on the day of purchase, showing a range of 26.1-79.8% human lineage in circulating cells (**Supplemental Table 2**). All mice in the treatment groups lost significant weight starting on day 3 after treatment but most mice regained weight starting on day 23 (**Figure 3A**). Two mice in the 111 kBq/g treatment group died of unknown causes, one on day 8 post injection and one on day 12. It is important to note that NSG mice (NOD/ltSz-scid Il2 rag -/-) contain mutations in *Prkdc* gene and other genes that make them extremely sensitive to radiation poisoning ^11^. Acute deaths were not seen in any nude mice given equal doses (**Figure 2 and Supplemental Figure 2**). Surviving mice were bled via mandibular vein on day 13-18 and again on day 28-31 and CBCs were obtained. [^212^Pb]-pent treatment caused a significant decrease in red blood cells, hemoglobin, and platelets from day 13-18, which partially recovered by day 28-31 (**Figure 3B**). No statistical changes were seen in WBCs on days 13-18; however, a significant leukocytosis was observed on days 28-31 in the 111 kBq/g treatment group (**Figure 3B**).

While hCD34+ humanized mice carry functional human bone marrow, they also maintain murine hematopoiesis. To distinguish damage to human bone marrow from murine bone marrow, the animals were sacrificed on day 28-31 and bone marrow was harvested from the femurs and tibias. Flow cytometry was performed using the gating strategy in **Supplemental Figure 5**, demonstrating a highly significant decrease in the frequency of human hCD45 cells in the [^212^Pb]-pent treatment groups compared to the controls. After treatment, measurable populations of viable hCD45+ cells remained in all animals (**Figure 4A**). Two mice retained larger populations of human bone marrow, with a frequency of 21.9% hCD45 in one 111 kBq/g animal and 11.7% hCD45+ in one 37 kBq/g treatment animal. Murine hematopoietic cells effectively reconstituted the bone marrow 28-31 days after [^212^Pb]-pent treatment, with bone marrow consisting of >75% mCD45+ cells in all treatment animals (**Figure 4B**). Among hCD45+ expressing cells, there was no difference in the percentage of bone marrow T cells between the control and treatment groups (**Figure 4C**). When compared with control animals, there was a statistically significant decrease in percentage of B-cells and a significant elevation in the percentage of CD14 expressing monocytes/macrophages at only the highest dose among the hCD45+ populations (**Figure 4C**). These data are consistent with the reported high affinity of pentixather for the human isoform of CXCR4 and indicate significant decreases in human hematopoietic cells, relative to the mouse hematopoietic cells, treated with therapeutic doses of [^212^Pb]-pent.

## Discussion

Small cell lung cancer remains notoriously difficult to treat despite improvements in our understanding of cancer biology. Phase III trials in recent years have shown that the addition of immune checkpoint inhibitors to chemotherapy in extensive stage SCLC improves median overall survival ≈ 2-3 months ranging from 12-16 months. However, few sizable improvements to treatment have been made for decades ^12-15^. Clearly, there is a critical need for novel therapy options in SCLC.

PRRT has become an established treatment in some neuroendocrine tumors, with therapy historically targeting the somatostatin receptor 2 (SSTR2). However, SSTR2 targeting radiopharmaceuticals are not frequently used in higher grade lung neuroendocrine carcinomas (NECs), such as SCLC, due to variable receptor expression ^16-18^. CXCR4, in contrast to SSTR2, has been shown to be highly expressed in SCLC ^3, 19^.

CXCR4 is a G-protein coupled receptor involved in hematopoietic stem cell migration and differentiation in normal tissues ^20^. In SCLC, CXCR4 activation has previously been implicated in tumor metastasis and chemoresistance ^21^, and CXCR4 expression has been linked with poor prognosis in SCLC ^2, 3^. Theranostic approaches directed against this receptor could benefit patients with some of the most aggressive SCLCs. Multiple clinical trials are investigating CXCR4 targeted ^68^Ga-pentixafor for PET/CT imaging in hematological malignancies (NCT05255926, NCT05093335, NCT05364177, NCT04561492, NCT04504526, NCT06125028) and in neuroendocrine tumors (NCT03335670). However, the current study is the first that we are aware of to demonstrate the efficacy of [^203^Pb]-pent for SPECT/CT imaging in SCLC.

One advantage of this approach is that CXCR4 dependent tumor binding of [^203^Pb]-pent in SCLC xenografts not only allows for visualization of radionuclide uptake in organs and tumors via SPECT/CT, but also allows for the accurate determination of dosimetry necessary for designing maximally effective therapy protocols. These data support the overall conclusion that [^203^Pb]-pent is an effective elementally matched imaging surrogate for [^212^Pb]-pent PRRT.

While established clinical practice with PRRT utilizes β- emitters, the ^212^Pb decay chain includes α- decay. α- particles exhibit high LET densely ionizing radiation tracks which generate free radical damage to cytosolic structures as well as double stranded DNA breaks. This results in markedly increased cell killing when compared to lower LET β-particles^22, 23^. Because of these benefits, ^212^Pb based α-particle PRRT shows promise for improving clinical responses in ongoing clinical trials in neuroendocrine tumors (NCT05636618, NCT06148636, NCT05153772), melanoma (NCT05655312), breast neoplasms (NCT01384253), cervical cancer (NCT05283330), and prostate cancer (NCT05720130). In addition, the short path length of an α-particle in solid tissues is only 25–80 µm compared to up to 1 cm in β-particles ^22^, which results in less nonspecific irradiation to adjacent tissues. The combination of these factors makes α-particle therapy a promising option to increase PRRT efficacy, but also raises questions about increased damage to healthy tissues expressing the target receptor.

Our preclinical studies were the first to investigate [^212^Pb]-pent therapy in SCLC and have demonstrated markedly increased overall survival and delayed tumor growth in SCLC xenografts. The absence of cytopenias as detected by CBC in treated animals, along with the lack of changes in liver and kidney fuction markers on serum chemistries, support the hypothesis that in this model, there is minimal off target damage associated with α-particle therapy. When compared with previous studies from our lab using the β- emitting radioligand ^177^Lutetium-pentixather ^3^, [^212^Pb]-pent appears to exhibit improved anticancer efficacy with a similar side effect profile in mice.

Myelotoxicity remains a concern with [^212^Pb]-pent treatment in humans due to the high expression of CXCR4 on hematopoietic stem cells. Consistent with previous comparisons of binding affinity for human CXCR4 versus mouse CXCR4 ^4^, current studies showed significant toxicity of [^212^Pb]-pent human hematopoietic lineages in CD34+ humanized mice 28-31 days following exposure. However, human bone marrow repopulation in this study is limited by competition with murine bone marrow that clearly recovered more quickly. Due to pentixather’s higher affinity for human compared to mouse CXCR4, treatment likely introduced a selective pressure against the repopulation of hCD45+ cells. Additionally, murine cells inhabit and are naturally homed to their native bone marrow niche, likely providing an additional selective advantage.Continued presence of viable human cells after treatment make it difficult to conclude that human hematopoietic cells could not reconstitute the bone marrow after [^212^Pb]-pent in humans without conducting a clinical study. However, hematopoietic stem cell harvest for autologous transplant may be required for [^212^Pb]-pent in humans.

Our preclinical data support the hypothesis that [^212^Pb]-pent therapy can be highly efficacious in SCLC patients with dismal prognoses. Due to the highly aggressive nature of SCLC, the [^203/212^Pb]-pent theranostic pair represents a novel option with the potential to prolong life in terminally ill patients. Our results have demonstrated the preclinical efficacy of [^212^Pb]-pent and warrant further investigation in a clinical study.

## Disclaimers

N/A

## Acknowledgments

This work was supported in part by NIH grants CA243014, CA217797, CA244091, CA243693, CA270742, P50 CA174521, CA086862, and the Holden Cancer Center. Data presented herein were generated in the Human Immunology Core and Flow Cytometry Facility, Carver College of Medicine / Holden Comprehensive Cancer Center, University of Iowa.

## Key Points

### Question

Can [^212^Pb/^203^Pb]-pent be used safely and effectively for imaging and therapy in SCLC in xenograft mouse models?

## Pertinent Findings

Mouse SCLC xenograft models demonstrated that [^203^Pb]-pent can effectively be used for CT/SPECT imaging and dosimetry calculations in CXCR4 expressing tumors. [^212^Pb]-pent was shown to be effective at treating SCLC xenografts, with some hematologic toxicity in hCD45+ cells.

## Implications for Patient Care

[^203/212^Pb]-pent have been shown to be highly efficacious for the treatment of SCLC in a preclinical model and warrant continuing investigation in clinical studies.

