## Supplemental figures and methods for "Targeting CXCR4 with [^212^Pb/^203^Pb]-Pentixather Significantly Increases Overall Survival in Small Cell Lung Cancer"

### Targeting CXCR4 with [$^{212}$Pb/$^{203}$Pb]-Pentixather Significantly Increases Overall Survival in Small Cell Lung Cancer – Supplemental Data and methods file.

**Supplemental Table 1: Expression of CXCR4 in Cell Lines used in Figure 1.**

CXCR4 expression in three SCLC lung cancer cell lines (DMS53, DMS273, H69AR) and one NSCLC cell line (H292) were determined via immunostaining and flow cytometry as previously described in reference 3 (PMID: 37989124). Results are reported as the ratio of MFI to that of the negative control, NSCLC cell line H292.

| Expression of CXCR4 in SCLC | | |
|---|---|---|
| Cell Type | Cell Line | MFI Ratio |
| NCSLC | H292 | 1 ± 0.08 |
| SCLC | DMS53 | 4.8 ± 0.6 |
| SCLC | DMS273 | 44.5 ± 5.9 |
| SCLC | H69AR | 84.9 ± 24.2 |

**Supplemental Figure 1: $^{212}$Pb-pent therapy at the 111 kBq/g or 37 kBq/g significantly prolonged survival and delayed tumor growth without unacceptable toxicity in nude mice bearing SCLC xenografts**.

[$^{212}$Pb]-pent therapy significantly prolonged survival and delayed tumor growth without significant weight loss in nude mice bearing SCLC xenografts. Athymic nude mice bearing DMS273 xenografts were treated with [$^{212}$Pb]-pent (111 kBq/g body weight; n=10) or equivalent volume of saline control (n=10) **A), B), C)**; or 37 kBq/g $^{212}$Pb-pent (n=9) or equivalent volume of saline control (n=9) **D), E), F).** Tumor growth was monitored via vernier calipers and is plotted. A significant delay in tumor growth was observed in the [$^{212}$Pb]-pent treatment groups ($p<0.05$). Mice were euthanized when tumor volume exceeded 1000 $mm^3$ twice. Kaplan Meier survival analysis demonstrated a significant increase in median overall survival in the $^{212}$Pb-pent treatment groups ($p<0.05$). No significant weight loss of other outward signs of distress were noted with these treatments.

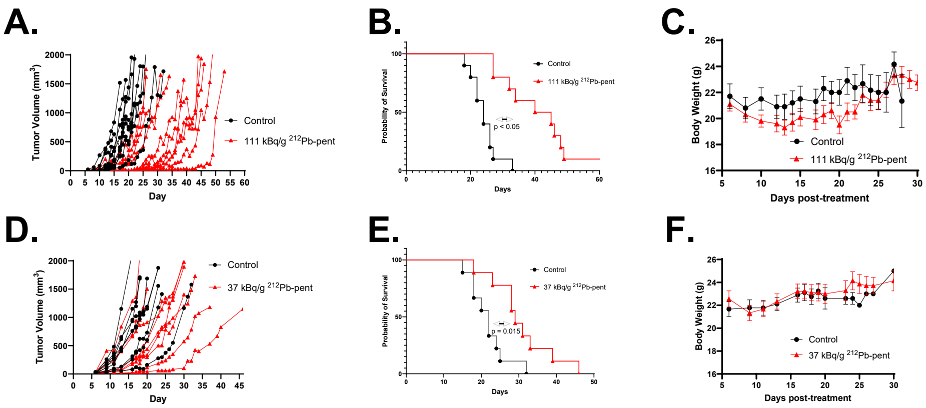


**Supplemental Figure 2: $^{212}$Pb-pent prolongs survival in mice bearing multi-drug resistant H69AR SCLC xenografts.** Athymic nude mice bearing high CXCR4 expressing, H69AR xenografts were injected with 55 kBq [$^{212}$Pb]-pent (n=4) or saline control (n=4) via tail vein injection. Animals were sacrificed when tumor volume exceeded 1000 $mm^3$ twice, and a significant survival advantage was observed in the [$^{212}$Pb]-pent treatment group with 3 of 4 animals showing no evidence of disease at 135 days.

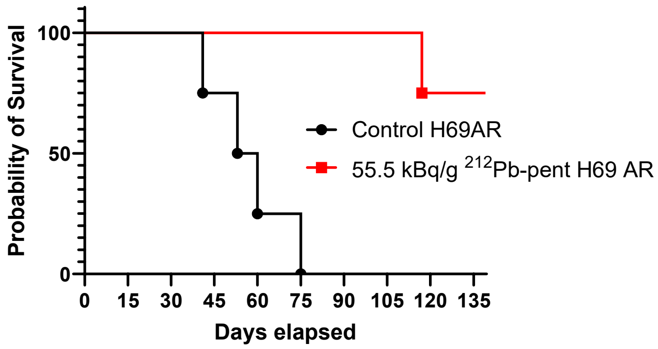

**Supplemental Figure 3: Gating scheme for analyzing long term hematopoietic stem cells (LT-HSC) with flow cytometry for nude mouse xenografts in main Figure 2D.**

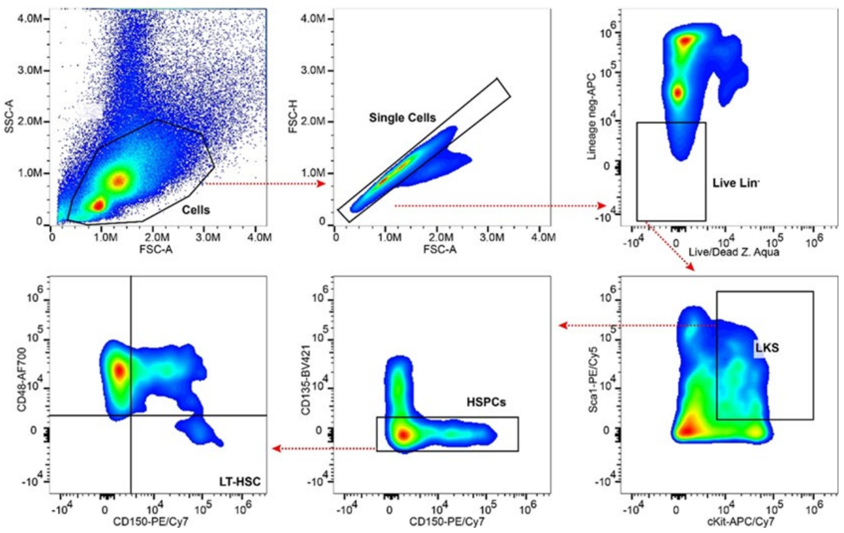

**Supplemental Figure 4: Mouse kidneys and livers show no biologically significant evidence of injury following [$^{212}$Pb]-pent exposure as determined by serum chemistries.** Blood was drawn from nude mice treated with saline control or 111 kBq/g [$^{212}$Pb]-Pent on the day of euthanasia from 5 randomly selected mice from the experiment shown in **Supplemental Figure 1A**. Serum chemistries were obtained and plotted for **A)** total serum protein, **B)** serum alanine aminotransferase **(**ALT), **C)** alkaline phosphatase (ALK Phos), **D)** total bilirubin, **E)** blood urea nitrogen (BUN), and **F)** creatinine. Dotted lines are normal published values in nude mice.

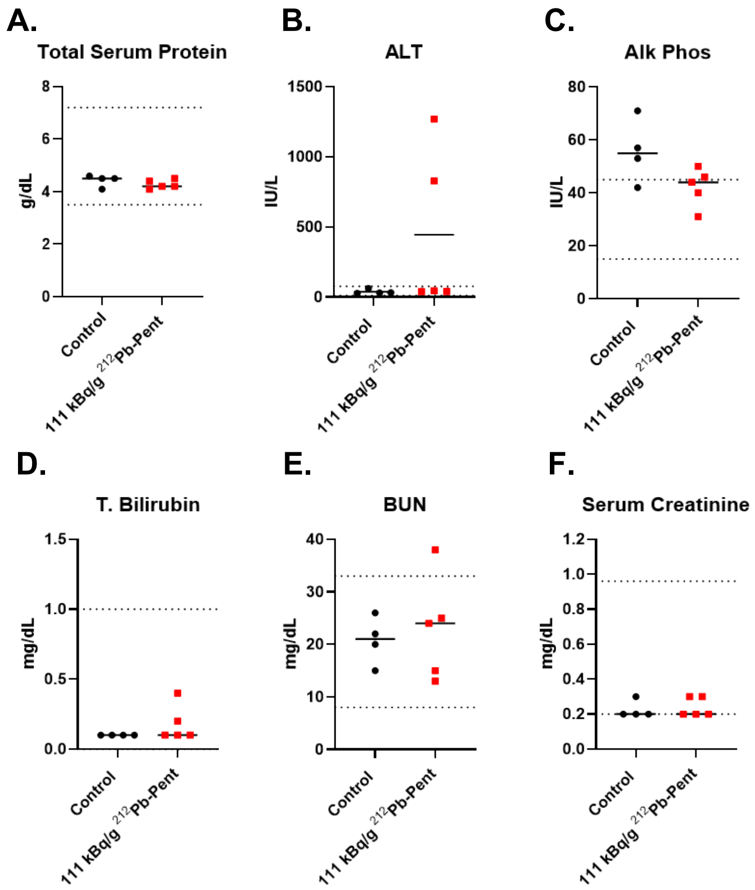


**Methods for Supplemental Figure 4:** Euthanasia was performed under analgesia with 200 µL intraperitoneal injection of ketamine xylazine 17.5/2.5 µg/µL. Adequate analgesia was determined by lack of toe pinch response, and blood was subsequently withdrawn via cardiac puncture followed by cervical dislocation. Samples were allowed to clot for 30 minutes and centrifuged at 8000 x g for 2 minutes. Serum samples were then submitted to Antech Diagnostic for analysis.

**Supplemental Table 2: Vendor provided analysis of peripheral blood cells from Humanized CD34+ mice purchased from Jackson Laboratory (Jackson 705557 Hu-NSG-CD34) and generated from NOD/SCID/IL2rγnull mice injected with human CD34+ hematopoietic stem cells. These animals were used in the experiments shown Figures 3 and 4.**

| Donor | hCD45+ | (B cell) | (T cells) | (myeloid) | mCD45+ |
|---|---|---|---|---|---|
| | Total % | % of hCD45 | % of hCD45 | % of hCD45 | Total % |
| 1245 hu NSG | 41.2 | 82.4 | 9.0 | 5.0 | 58.8 |
| 1245 hu NSG | 33.6 | 88.6 | 4.0 | 4.5 | 66.4 |
| 1245 hu NSG | 55.6 | 87.7 | 5.0 | 4.0 | 44.4 |
| 1246 hu NSG | 79.8 | 84.4 | 1.0 | 4.9 | 20.2 |
| 1246 hu NSG | 73.9 | 85.0 | 0.0 | 5.3 | 26.1 |
| 1246 hu NSG | 75.0 | 75.2 | 14.0 | 3.1 | 25.0 |
| 1247 hu NSG | 26.1 | 49.4 | 31.9 | 9.1 | 73.9 |
| 1247 hu NSG | 33.5 | 64.6 | 14.7 | 9.7 | 66.5 |
| 1247 hu NSG | 36.4 | 79.1 | 1.5 | 10.2 | 63.6 |
| 1753 hu NSG | 39.8 | 72.2 | 21.7 | 2.4 | 60.2 |
| 1753 hu NSG | 30.5 | 72.3 | 21.4 | 2.5 | 69.5 |
| 1753 hu NSG | 68.5 | 83.5 | 8.0 | 5.2 | 31.5 |
| 1754 hu NSG | 43.7 | 89.7 | 0.8 | 1.8 | 56.3 |
| 1754 hu NSG | 46.3 | 85.1 | 3.1 | 3 | 53.7 |
| 1754 hu NSG | 69.1 | 87.7 | 0.9 | 3.2 | 30.9 |
| 1729 hu NSG | 39.4 | 89.8 | 0.3 | 3.8 | 60.6 |
| 1729 hu NSG | 52.1 | 85.7 | 3.1 | 5.5 | 47.9 |
| 1729 hu NSG | 50.2 | 86.2 | 1.9 | 5.1 | 49.8 |

**Supplemental Figure 5: Gating strategy for flow cytometry of CD34+ humanized mouse bone marrow in Figure 4**

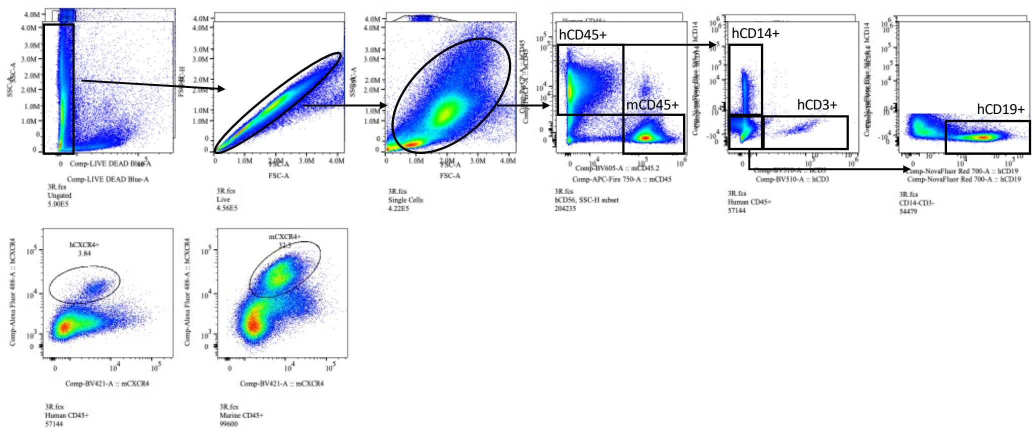
